# Theta rhythm supports hippocampus-dependent integrative encoding in schematic memory networks

**DOI:** 10.1101/2019.12.16.874024

**Authors:** Berta Nicolás, Jacint Sala-Padró, David Cucurell, Mila Santurino, Mercè Falip, Lluís Fuentemilla

## Abstract

Integrating new information into existing schematic structures of knowledge is the basis of learning in our everyday life activity as it enables structured representation of information and goal-directed behaviour in an ever-changing environment. However, how schematic mnemonic structures aid the integration of novel elements remains poorly understood. Here, we showed that the ability to integrate novel picture information into learn structures of picture associations that overlap by the same picture scene (associative network) or by the conceptually related scene information (schematic network) is hippocampus-dependent, as patients with lesions at the medial temporal lobe (including the hippocampus) were impaired in inferring novel relations between elements within these mnemonic networks but not in retrieving individual pictures in a subsequent memory test. In addition, we observed more persistent and widespread scalp electroencephalographic (EEG) theta oscillatory pattern (3-6Hz) while healthy participants encoded novel pictures related to schematic memory networks, suggesting that theta may reflect distances between elements within a representational network space. Finally, we found high similarity values for neural activity patterns elicited by novel and related events only within associative networks, thereby suggesting that neural reactivation may promote the integration of new information into existing memory networks only when direct associations within the network link their elements. These findings have important implications for our understanding of the neural mechanisms that support the development and organization of structures of knowledge.

## Introduction

Experiences often overlap in content, presenting opportunities to integrate them into mnemonic networks. These mnemonic networks share certain characteristics, such as plasticity and hierarchical or schematic organisation (Eichenbaum 2017), which enable structured representation of information and goal-directed behaviour in an ever-changing environment (McKenzie et al. 2014). However, how such schematic mnemonic structures aid the integration of new information remains unclear.

The standard approach to examining integrative encoding into memory networks has been to train subjects on separate events that share common elements (e.g., AB and BC) and then test for the associative network (ABC) via assessment of knowledge about the indirectly associated network elements (AC). This research has shown that the hippocampus and the prefrontal cortex (PFC) are not essential to training on individual associations (AB and BC) but do play a critical role in integrating information across related associated events (AC) (Dusek and Eichenbaum 1997; Greene et al. 2006; Heckers et al. 2004; Preston et al. 2004; Schlichting and Preston 2016). Leveraged by the use of neuroimaging techniques with fine-temporal resolution, such as magnetoencephalography (MEG) and electroencephalography (EEG), recent studies have also shown that the integration of novel events into an existing associative memory network relies on hippocampus-driven oscillatory activity in the theta range (3-8Hz) (Backus et al. 2016; Sans-Dublanc et al. 2017). Thus, while this approach has provided valuable insights into the neural underpinnings supporting the formation of associative memory networks, we still lack understanding of whether a similar neural framework can be generalized to more complex scenarios, akin to real-life environments, whereby schematic structures of knowledge foster the rapid assimilation of new events (Van Kesteren et al. 2010, 2012; Packard et al. 2017; Tse et al. 2011);

To address this issue, we designed a two-phase task wherein participants learned an intermixed set of picture associations (i.e., face-scene) that overlapped with a common scene (i.e., associative network condition) or with scene images that depicted the same conceptual information (i.e., schematic network condition) and subsequently generalized to novel stimulus combinations (Figure 1). During learning phase 1 (LP1), participants learned to associate a face with a scene by choosing which of two scenes went with the face, and then receiving feedback. While each face-scene association was learned individually, there was partial overlap across events. Some pairs overlapped with a common face picture (A-B and A-C; associative network condition) and other pairs with scene pictures from the same semantic category (A-B_1_ and C-B_2_; schematic network condition) (Figure 1A). By the end of LP1, we expected that participants would have successfully learned the individual associations and integrated them based on their relational network properties, namely associative or schematic (Figure 1D).

**Figure 1.**
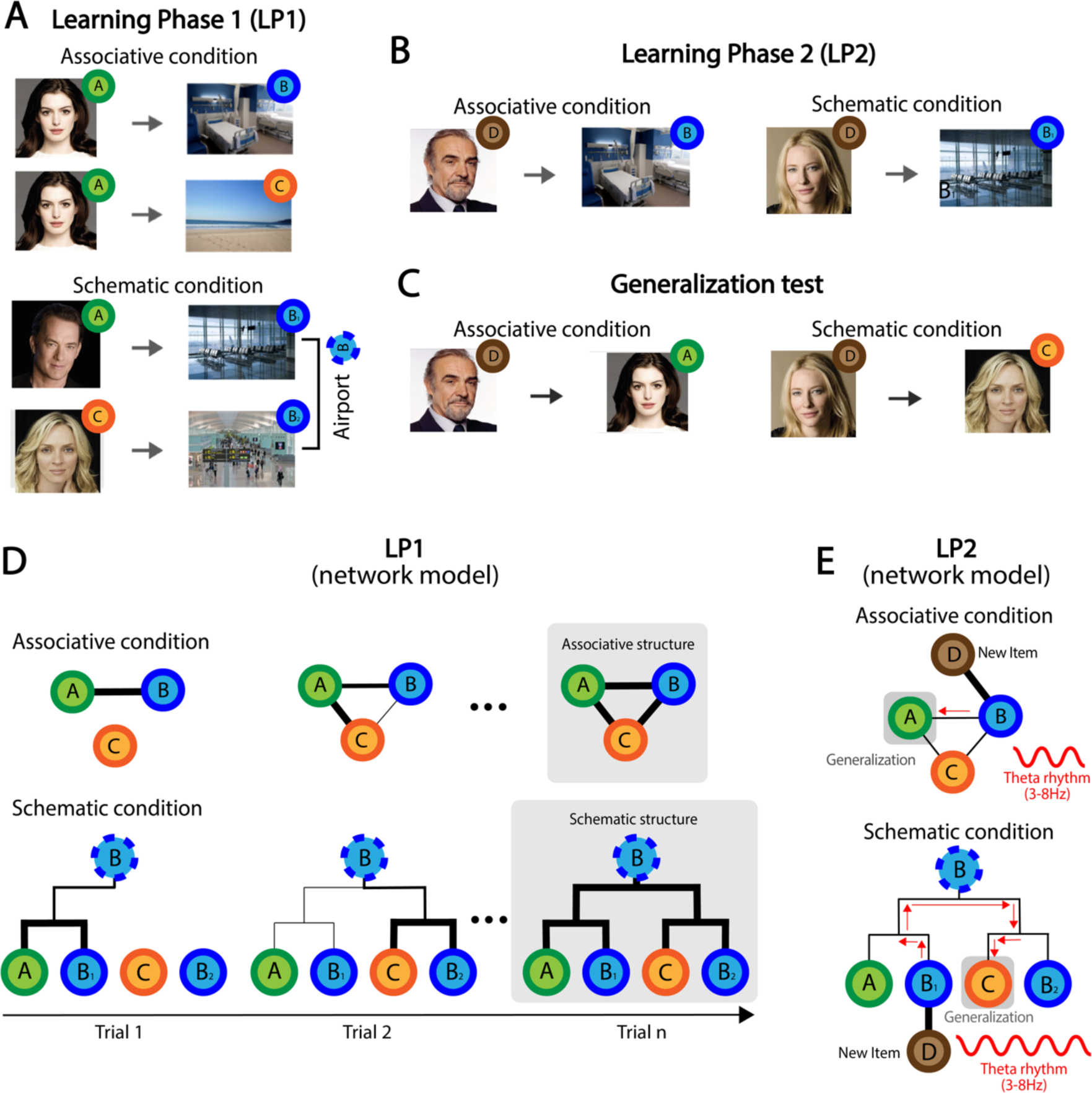
Experimental design and mnemonic network models. **(A)** In LP1, participants encoded pairs of face-scene images. Picture pairs were organized so that some of them shared the face image (Associative condition) and some shared the scene semantic context (Schematic condition). Picture pair conditions were presented intermixed during LP1. (**B**) In a following LP2, participants had to learn novel face-scene pairs. All pairs used face images that overlapped with one of the pair images from each subset in LP1. Picture pair conditions were presented intermixed during LP2. (**C**) Participants’ memory for LP1 and LP2 picture pairs was subsequently tested using a two-alternative forced choice paradigm that included directly learned association trials (“trained”) as well as inference trials that tested participants’ ability to generalize. Specifically, “generalization” trials tested whether participants would choose A/C, encoded in LP1, when presented D, encoded in LP2. (**D**) Hypothesized memory representation model accounting for each of the learned picture sets in the associative and schematic conditions throughout LP1. Thick lines between elements depict picture pairs presented in a given trial. Thin lines depict connections of picture pairs established during learning via integrative encoding. At the end of LP1, we hypothesized, several corresponding memory networks were acquired, and their structure reflected an associative or a schematic typology. (**E**) Diagram depicting our hypothesis that the integration, during LP2, of a novel element in the establish memory networks from LP1 promoted the binding of a connected set of nodes within the network and that this process was signalled by theta oscillations. A and B depict face images of famous people due to bioRxiv policy on not displaying pictures of real people. Face images used in our study were taken from Minear et al. (2004), which provide a face image database with the authorization for publication for research purposes.

To assess whether these two mnemonic network structures influenced integrative encoding of new information, we next asked participants to encode a novel set of face-scene picture associations (i.e., learning phase 2, LP2), wherein each scene picture corresponded to a scene from the mnemonic picture set learned in LP1 (Figure 1B). Thus, we expected that the overlap between scene images (B and B_1_ in Figure 1B) would induce the integration of the new face images (i.e., D) into the specific memory network learned in LP1, thereby promoting generalization (Figure 1E). After LP2, participants were tested using a two-alternative forced choice paradigm that included directly learned association trials (“trained”) as well as inference trials that tested participants’ ability to generalize. Specifically, “generalization” trials tested whether participants would choose A/C, encoded in LP1, when presented D, encoded in LP2 (Figure 1C), thereby assessing whether they had successfully integrated LP2 events into the related memory structures acquired in LP1.

Here, we aimed at examining whether events encoded in LP2 elicited different theta oscillatory patterns as a function of whether they were linked to associative or schematic memory networks acquired in LP1. Previous findings have shown that theta activity encoded representational distances in a spatial space (Bush et al. 2017; Vass et al. 2016) but also in the semantic and temporal word space (Solomon et al. 2019), supporting the idea that theta underlies the navigation through a general-domain cognitive map in the hippocampus. Thus, we hypothesized that representational distance between the new and the rest of the items within the schematic memory network was much greater than the distances within an associative memory network, and that this would be reflected as more persistent theta activity elicited by each of the events in LP2 (Figure 1E). Second, prior literature has emphasized that inferential learning relies on the reactivation of memory events related to a network (e.g., (Zeithamova et al., 2012)). In the current study, we implemented a time-resolved neural similarity analysis (e.g., Silva, Baldassano, and Fuentemilla 2019; Sols et al. 2017) to elucidate whether LP2 events elicited the reactivation of elements within a schematic and associative network. And third, we examined the critical role of the hippocampus in integrative encoding for associative and schematic memory networks by comparing behavioural data from chronic epileptic patients with lesion at the hippocampus with data from a matched control sample.

## Material and Methods

### Participants

#### Experiment 1: Healthy adults

Forty right-handed healthy volunteers (34 females) participated in the experiment 1. Mean age of the participants was 23.62 (SD = 2.98 years). All the participants included in the study reported no history of medical, neurological, or psychiatric disorders, and no drug consumption. All subjects were volunteers, gave written informed consent, consented to publication, and received financial compensation for their participation in this study. All participants had normal or corrected-to-normal vision. The study was approved by the Ethics Committee of the University of Barcelona.

#### Experiment 2: TLE patients

A group of fourteen patients with refractory mesial temporal lobe epilepsy (TLE) caused by different aetiologies was recruited following a pre-surgical evaluation at the University Hospital of Bellvitge (Table 1). All patients had sustained damage to the right or left anterior medial temporal lobe structures, including the hippocampus. In all patients, verbal and non-verbal intelligence was assessed using the Weschler Memory Scale, and the mean IQ was 92.30 (±10.72). None of them showed mental disabilities (IQ under 80). Patient diagnosis was established according to clinical, EEG, and magnetic resonance imaging or FDG18PET. All of the patients underwent neurological and neuropsychological examination, continuous video-EEG monitoring, and structural and functional neuroimaging (MRI and PET). Patients were included in the study when the clinical data, EEG findings, and neuroimaging data suggested unilateral mesial TLE. All patients had: 1) seizures with typical temporal lobe semiology that were not controlled with antiepileptic drugs, 2) EEG patterns concordant with mesial temporal lobe epilepsy, and 3) neuroimaging data supportive of hippocampal involvement in seizure generation. None of the patients suffered a seizure during the experimental task or 24 hours before the task, and all of the patients were on habitual anti-epileptic drug regimens. The study was approved by the Ethical Committee of University Hospital of Bellvitge. Informed consent was obtained from all of the patients before participation in the study.

**Table 1.**
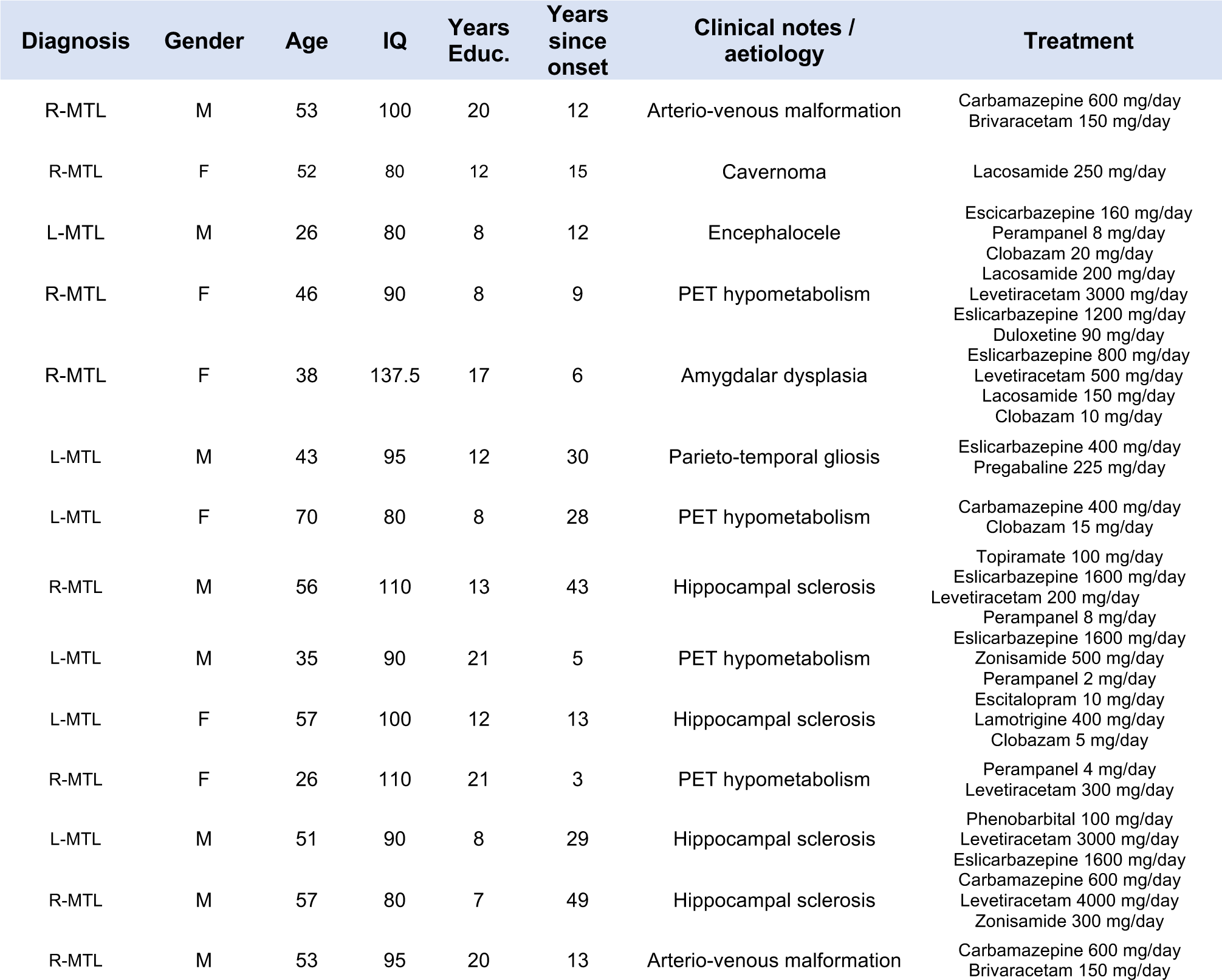
Individual patient characteristics.

#### Experiment 2: Healthy controls

The control group consisted of fourteen participants with no history of neurological disorders. Control participants were individually matched to TLE patients. No differences were found between groups in terms of age (t (13) = −1.794, p = 0.096) or years of education (t (13) =1.325, p = 0.208). Informed consent was obtained from all subjects before their participation in the study.

### Experimental procedures

#### Stimuli

Stimuli consisted of 24 images of Caucasian (half female and half male) non-expressive faces (F) from Radboud database (Langner et al. 2010) and from UT Dallas database (Minear and Park 2004) and 48 scene (S) images selected to depict 48 real-life contexts, half of them from indoor contexts (bars, airports, hospitals, supermarkets, kitchens, bakeries, hairdresser’s, clothing stores, locker room, cinema, bus stop, ice-cream shop, computer store, campsite, jail) and the other half from outdoor contexts (parks, landscapes, waterfalls, mountains, caves, beaches, lakes, forests), from SUN database (Xiao et al. 2010).

The faces were distributed through LP1 (8 for the episodic and 16 for the semantic experimental condition), LP2 (8 for each experimental condition), generalization (8 for each experimental condition), and trained test (16 each experimental condition). Scene context images were distributed over LP1 (16 for each experimental condition) and were repeated over LP2 (8 for each experimental condition) and trained test (16 for each experimental condition). F-S pair assignments were randomized between participants. The order of F-S picture presentation was randomized within each LP1 and LP2 block.

#### Task design and procedure in healthy participants (Experiment 1)

Participants performed a modified version of an associative inference task (e.g., (Zeithamova et al., 2012) (Figure 1A). The task consisted of two separate learning phases, Learning Phase 1 (LP1) and Learning Phase 2 (LP2), followed by a Testing Phase. In each of the acquisition phases, participants were requested to learn Face (F) – Scene (S) associations through feedback. Unbeknownst to the participants, in LP1 F-S associations were organized into eight picture subsets, each including two F-S pairs. Subsets in the associative condition involved one face image (F1) and two different scene images (S1 and S2). During learning, F1 was presented separately from S1 and S2, thereby promoting inferential learning between S1 and S2 through associative overlap. Picture subsets in the semantic condition involved two different face images (F1 and F2) and two different scene images (S1 and S2) from the same semantic category. Specific F-S pairs in the semantic condition were randomly assigned before the experiment started. LP1 was structured into 8 blocks, each including 32 trials in total (16 trials per experimental condition). LP1 was followed by LP2, which consisted of 16 different F-S pairs presented 8 times throughout the 8 blocks. Importantly, in LP2 all face images were novel but each of them was paired with a scene image from a different subset of pictures presented in LP1. Thus, 8 faces were associated with 8 scenes from each of the associative condition subsets (henceforth, associative condition in LP2) and 8 faces were presented with 8 scenes from each of the schematic condition subsets (semantic condition in LP2, thereby providing opportunities for integrative encoding of picture subsets from LP1 that had differential relational structure (i.e., associative or schematic)).

The structure of the trials was similar in LP1 and LP2. Each trial consisted of the presentation of a face at the top of the screen and two scenes at the bottom for 3500 ms. Participants had to wait for the appearance of the message ‘RESPONSE’ and they then had 1000 ms to indicate, by pressing a button, which of the two scenes was associated with the face. Following participants’ choices, a delay period (grey background) of 500 ms preceded the feedback, which consisted of the presentation of a green tick (right choice) or a red X (wrong choice), each of which remained on the centre of the screen for 1000 ms. The appearance of the scenes on the right or left side of the screen was counterbalanced through the presentations. Additionally, in order to avoid stimulus-response learning strategies, every scene was shown as a correct choice for a particular face and as an incorrect choice when appearing with other faces, with the restriction that it could not appear twice as an incorrect choice with the same face. Therefore, the correct scene for a given face was always the same, but the incorrect scene was variable.

Two separate surprise force-choice tests followed LP2: the generalization and the trained test. In the generalization test, participants had to indicate which of two faces seen during LP1 was associated with a face from LP2 presented at the top of the screen, thereby assessing inferential learning. This test consisted of 16 trials (8 for each experimental condition). In the trained test, two scene images appeared, and participants had to indicate which had been associated with the face image presented at the top of the screen. This test always followed the generalization test, thereby ruling out the possibility that accuracy in the inferential test could be explained by other factors (i.e., memory recall for pair associates) other than neural mechanisms elicited during LP2. In this test, all trained pairs from LP1 and LP2 were tested. In the generalization and direct test, the incorrect choice elements were all previously learned items that had been studied during the task. Pictures remained on screen until the participants responded, and there was no feedback informing the participants of the result of their choice. Test trials were separated by an inter-trial time randomized between 750 and 1250 ms.

#### Task design and procedure in TLE patients and healthy controls (Experiment 2)

A shorter version of the experiment 1 task was implemented in TLE patients and control samples. More concretely, LP1 consisted of 12 subsets of F-S associations: 6 from the associative condition and 6 from the schematic condition. All pairs were presented 8 times throughout 8 different blocks. LP2 consisted of 6 face-context associations (3 from the associative condition and 3 from the schematic condition). At the end of the task, a generalization test consisting of 6 possible face-face associations, and a trained test consisting of 12 possible face-context associations were implemented.

#### Behavioural analysis

Participants’ correct responses throughout LP1 and LP2 were calculated and averaged for each block of trials. A repeated measures ANOVA including Block (8 levels) and experimental condition (associative and schematic) as within-subject factors was used for statistical assessment. Participants’ accuracy in the tests was assessed by the proportion of correct choices separately for the generalization test and the trained test. Statistical significance was set at an alpha of 0.05. Greenhouse-Geisser epsilon correction was used to correct for possible violations of the sphericity assumption for statistical analysis when necessary; the adjusted p-values after the correction were reported.

#### EEG recordings and preprocessing in Experiment 1

EEG was recorded at a 500 Hz sampling rate (High-pass filter 0.01Hz, notch filter at 50Hz) from the scalp using a BrainAmp amplifier and tin electrodes mounted on an electrocap (Electro-Cap International) located at 29 standard positions (Fp1/2, Fz, F7/8, F3/4, FCz, FC1/2, FC5/6, Cz, C3/4, T3/4, Cp1/2, Cp5/6, Pz, P3/4, T5/6, PO1/2, Oz) and at the left and right mastoids. An electrode placed at the lateral outer canthus of the right eye served as an online reference. EEG was re-referenced offline to the linked mastoids. Vertical eye movements were monitored with an electrode at the infraorbital ridge of the right eye (EOG channel). Electrode impedances were kept below 3 kΩ. EEG was band-pass filtered offline at 0.1 – 40Hz. Independent Component was applied to the continuous EEG data to remove blinks and eye movement artefacts. Trials exceeding ± 100 μV in both EEG and EOG within a −100 to 2500 ms time window from stimulus onset were rejected offline and not used in the time-frequency and neural similarity analysis detailed below. 7 participants were excluded from subsequent EEG analyses as they did not produce at least 5 artefact-free trials for each of the 8 learning blocks in LP2.

#### Time-frequency (TF) analysis

TF was performed using six-cycle complex Morlet wavelets in 7100 ms EEG epochs (2100 ms before stimulus onset through 5000 ms after) from LP2. Changes in time-varying energy (square of the convolution between wavelet and signal) in the 2-14 Hz band were computed for each trial and averaged separately for each experimental condition at the individual level. Before performing an overall average, power activity changes were computed with respect to the baseline of each participant (−200 to 0 ms from picture onset).

#### Similarity analysis

This analysis was set to assess for the possibility that the encoding of LP2 picture pairs elicited the reactivation of neural patterns triggered by picture pairs from the same memory network acquired in LP1, thereby suggesting, according to previous reports investigating inferential learning (Zeithamova et al., 2012), that neural reactivation arises as a mechanism supporting integrative encoding during LP2. To address this issue, we implemented a time-resolved trial-to-trial similarity analysis between EEG patterns elicited during the last block in LP1 and EEG patterns elicited throughout LP2. We reasoned that including trials only from the last LP1 block guaranteed that neural patterns taken in the analysis were the strongest and most stable memory traces associated with each picture pair in LP1, as learning accuracy in that block was almost perfect and similar between experimental conditions (see results below).

The similarity analysis was performed at the individual level, and included spatial (i.e., scalp voltages from all the 29 electrodes) and temporal features, which were selected in steps of 10 sample points (20 ms) of the resulting z-transformed EEG single-trials. Similarity analysis was implemented at single-trial level by correlating point-to-point the spatial EEG features throughout 2500 ms from picture onset. The similarity analysis was calculated using Pearson correlation coefficients, which are insensitive to the absolute amplitude and variance of the EEG response. R values were then Fischer z scored before statistical comparison analysis.

#### Cluster statistics of the EEG data

To assess for power differences between conditions at the temporal domain, we used a paired sample permutation test (Groppe, Urbach, and Kutas 2011) to deal with the multiple comparisons problem given the multiple sample points included in the analysis. This test uses the “*t* max” method to adjust the p-values of each variable for multiple comparisons (Blair and Karniski, 1993). Like Bonferroni correction, this method adjusts p-values in a way that controls for the family-wise error rate.

To account for scalp distribution differences between associative and schematic conditions in time-frequency data and to account for differences between conditions in the similarity analysis, a cluster-based permutation test was used (Maris and Oostenveld 2007), which identifies clusters of significant points in the resulting 2D matrix in a data-driven manner and addresses the multiple-comparison problem by employing a nonparametric statistical method based on cluster-level randomization testing to control for the family-wise error rate. Statistics were computed for each time point, and the time points whose statistical values were larger than a threshold (p < 0.05, two-tail) were selected and clustered into connected sets on the basis of x,y adjacency in the 2D matrix. The observed cluster-level statistics were calculated by taking the sum of the statistical values within a cluster. Then, condition labels were permuted 1000 times to simulate the null hypothesis, and the maximum cluster statistic was chosen to construct a distribution of the cluster-level statistics under the null hypothesis. The nonparametric statistical test was obtained by calculating the proportion of randomized test statistics that exceeded the observed cluster level statistics.

## Results

### Experiment 1 (Healthy participants)

#### Behavioural performance

All participants were able to learn face-scene associations from the associative and schematic condition in LP1 (Figure 2A). This was reflected by high accuracy (i.e., > 90%) in the participants’ ability to choose the association pair correctly in the two conditions in the last block of the encoding paired t-test: t(39) = 1.50; p = 0.14). A repeated-measures ANOVA including condition (associative and schematic) and block (from one to eight) as within-subject factors confirmed accuracy improvement over the course of the task for all subsets of pictures (main effect of block: F(4.03,157.22) = 174.96, p < 0.01). However, that increment was less steep in the schematic than in the associative condition (Condition × block effect: F(5.01,195.47) = 2.51, p = 0.016).

**Figure 2.**
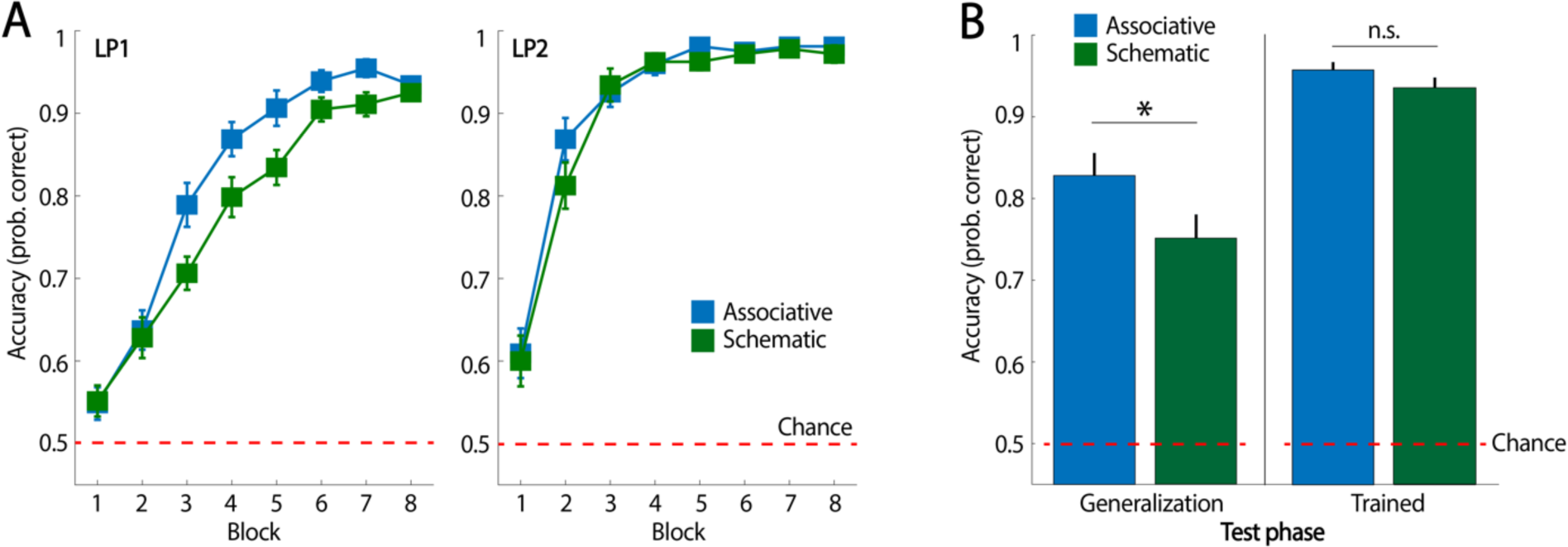
Behavioural data in healthy young participants (Experiment 1). **(A)** Averaged participants’ accuracy in selecting the correct scene association with a given face throughout LP1 and LP2 for each experimental condition. (**B**) Averaged participants’ accuracy in the generalization and the trained memory tests. *p < 0.05 and n.s., p > 0.05. Error bars indicate standard error of the mean.

In LP2, participants reached high levels of accuracy relatively rapidly and they were highly accurate (i.e., > 90%) in selecting the correct association by the end of the learning phase (Figure 2A). A repeated-measures ANOVA, including experimental condition and block as within-subject factors, revealed no significant differences between conditions (F(1,39) = 2.56, p = 0.12) or experimental condition × block (F(7,273) = 0.78, p = 0.60). A trend towards significance was found for the block factor (F(5.20,202.99) = 1.87, p = 0.07), indicating that participants’ learning occurred very rapidly during encoding and reached a ceiling effect at early stages of the encoding rounds.

Participants showed, overall, high accuracy in the generalization test, thereby demonstrating that they had successfully integrated picture sets from LP1 during LP2 (Figure 2B). However, we found that accuracy was greater in the associative (Mean = 82.81%, SD = 17.15%) than the schematic condition (Mean = 75%, SD = 18.95%) (t(39) = 2.48, p = 0.018). Importantly, participants were highly accurate in the trained test (i.e., > 80%) and their performance did not differ between learnings and conditions (F(1,39) = 0.12, p = 0.73) (Figure 2C), thereby confirming that they retained the trained associations from the two experimental conditions in equal measure. Altogether, the behavioural findings suggest that the underlying structure of the associations learned during LP1 may have had an impact during encoding strategies in LP2, which is when participants had the possibility to create the relational links needed to establish inferential learning between face-scene pairs.

#### Theta oscillations

To test our hypothesis that theta oscillations would persist longer in time in response to schematic than associative conditions, we performed a time-resolved comparison of theta power changes elicited during the 2500 ms after stimulus onset between associative and schematic conditions. We targeted a broad frequency range of theta oscillations spanning 3 – 6 Hz, which is a slightly lower frequency than the traditional theta band according to recent reports (Jacobs et al. 2013; Watrous et al. 2013), and to scalp electrophysiological findings using similar experimental designs (Backus et al. 2016; Sans-Dublanc et al. 2017). In line with this literature, theta power changes at this frequency range were pronounced during LP2 associative and schematic condition trials (Figure 3). Importantly, and confirming our hypothesis, we found that the theta power increase was more extended in time in response to picture pairs that were linked to schematic rather than to associative memory structures acquired in LP1 (Figure 3). A cluster-based permutation test revealed that the persistent theta power increase in the schematic condition as compared to the associative condition was distributed over the scalp, spanning frontal, central, and posterior scalp sensors (Figure 3).

**Figure 3.**
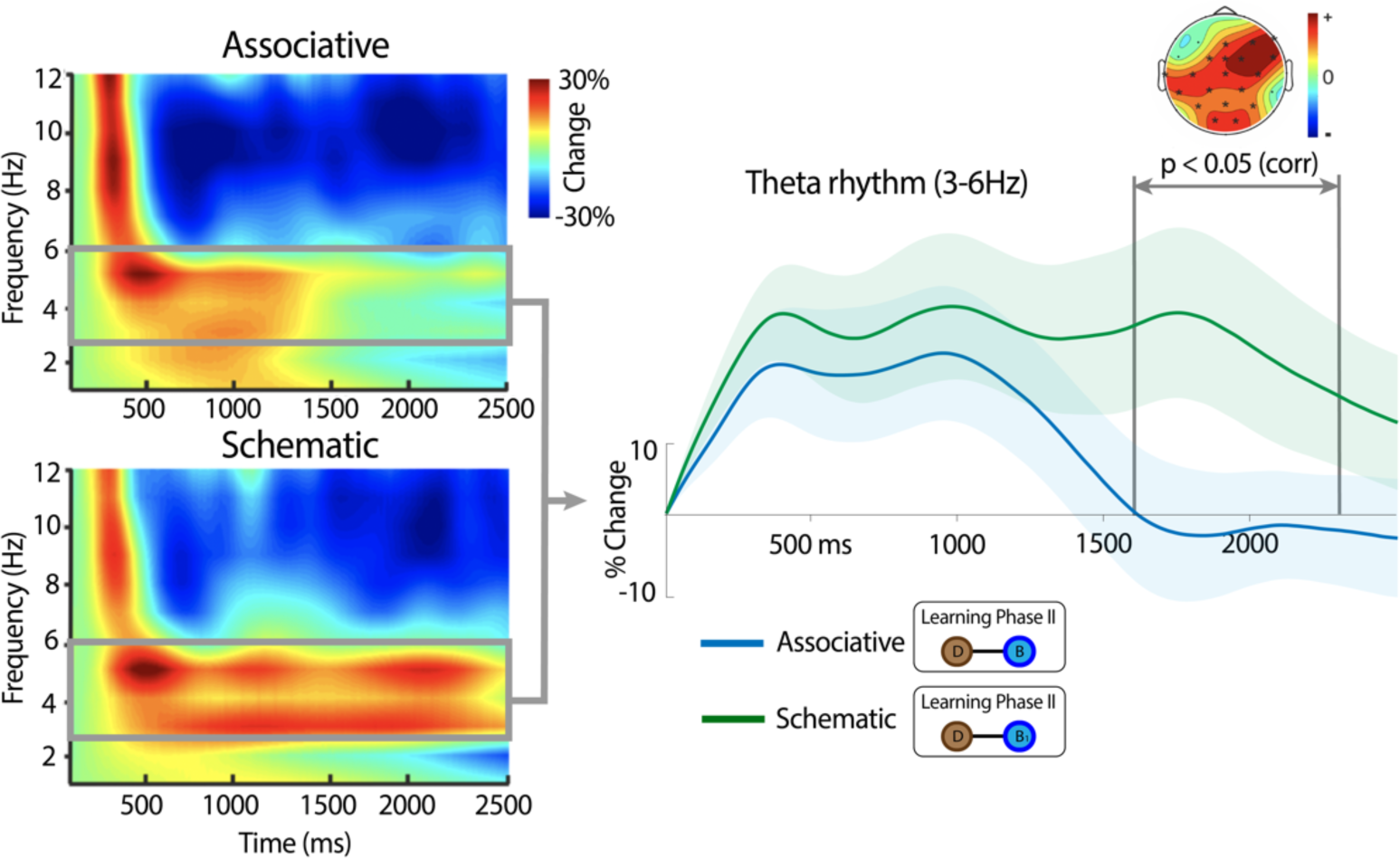
Theta oscillations in LP2. Group-averaged changes in spectral power (averaged over all scalp sensors) elicited by picture pairs from the associative and schematic conditions in LP2. A power increase in the theta band was observed in both conditions. However, that power increase lasted longer during the encoding of picture pairs from the schematic than the associative condition. Statistical time window differences between conditions are indicated with bars (point-to-point paired *t*-test threshold p < 0.05). Significant time points corrected for multiple comparison were between 1730 and 1950 (p < 0.05; one-tail) within that time window. Spatially distributed theta differences between schematic and associative conditions are also depicted; sensors that were significant and corrected for multiple comparisons at cluster level are marked with a black asterisk. Thick theta line represents the mean across participants and point-to-point standard error is depicted in shaded colour.

#### Neural similarity

This analysis revealed that the patterns of EEG responses elicited by picture pairs in the last LP1 block correlated with EEG patterns elicited during the encoding of different picture pairs that overlapped in content (Figure 4A). However, the degree of neural similarity differed between experimental conditions. More specifically, picture pairs linked to learned picture pairs in LP1 from the associative condition showed significantly stronger neural similarity values over a window of ~500 to 2000 ms from stimulus onset in LP2. On the other hand, EEG patterns elicited by picture pairs in the schematic condition showed an increased neural similarity earlier in time during the encoding of linked LP2 picture pairs (at around 300-500 ms from LP2 picture onset) (Figure 4B). In addition, the same analysis of LP1 trials from the first block during learning did not show any statistically significant differences between trial conditions, thereby suggesting that the similarity effects were greatest when neural patterns reflected robust memory representations of network representations at the end of LP1 (e.g., Figure 1D).

**Figure 4.**
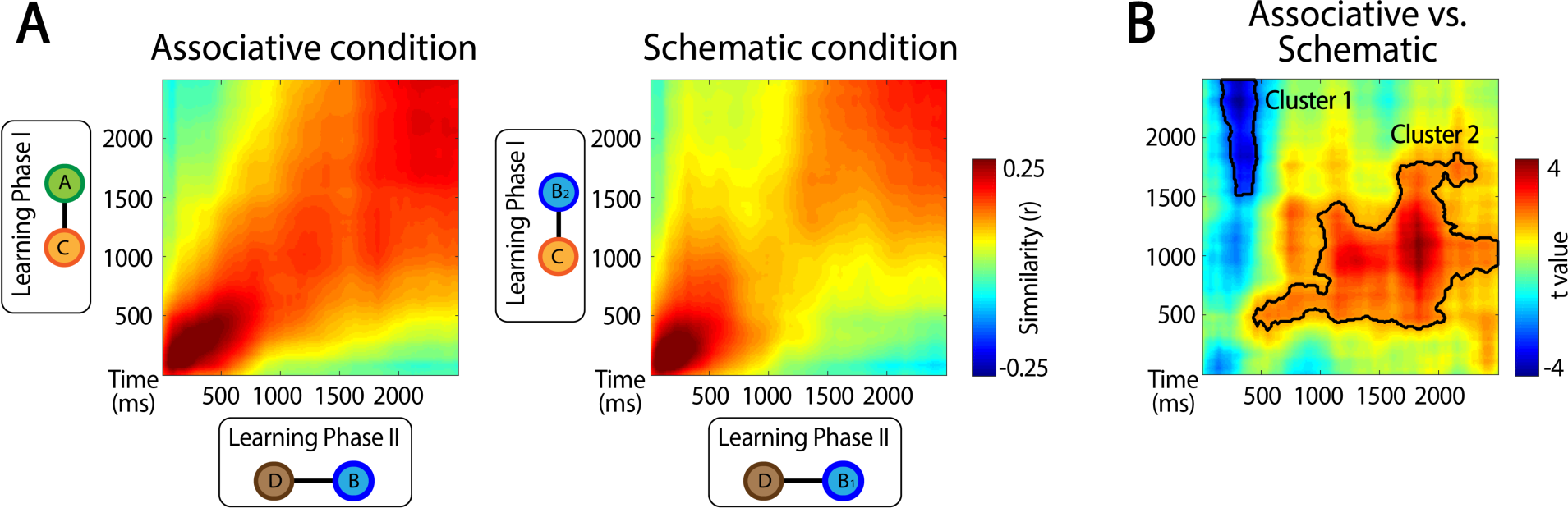
Neural pattern similarity between events that overlapped within memory network structures in the task. (**A**) Group-averaged time-resolved degree of similarity between C-A and C-B_1_ events from the last block of trials in LP1 and the corresponding D-B trials during LP2. Neural similarity in this analysis offers a measure of how similar two neural patterns are when elicited by events which, although separated in time, share partial memory information in the current task. (**B**) Point-to-point *t* value map from comparing schematic and associative neural similarity results. Two clusters of statistically significant similarity values were found (p < 0.05, cluster-based permutation test) (indicated by a thick black line).

### Experiment 2 (TLE patients and matched controls)

In line with experiment 1, a mixed-design ANOVA, including condition and block as a within-subject factor and group (TLE and control) as a between factor in LP1 data, revealed a statistically significant main effect in condition (F(1,26) = 5.14, p = 0.03) and block (F(7,182) = 19.35, p < 0.01), and a non-significant condition × block interaction (F(7,182) = 1.37, p = 0.22), which indicates that TLE patients and controls successfully encoded picture pairs over the task but that events from the associative condition were learned faster. No significant differences were found between groups in any of the contrasts (i.e., condition × group: F (1, 26) = 0.6, p = 0.81, block × group: F (7,182) = 1.66, p = 0.12; and condition × block × group: F (7,182) = 2.03, p = 0.053). The same statistical analysis of behavioural data from LP2 revealed that both groups successfully acquired picture pairs over the course of the task (block effect: F(4.13,107.37) = 15.64, p < 0.01; block × group interaction: F(7,182) = 0.80, p = 0.58), independently of the experimental conditions (condition effect: F (1,26) = 0.03, p = 0.87; condition × group interaction: F(1,26) = 1.11, p = 0.30); condition × block × group: F(7,182) = 0.95, p = 0.47)) (Figure 5A).

**Figure 5.**
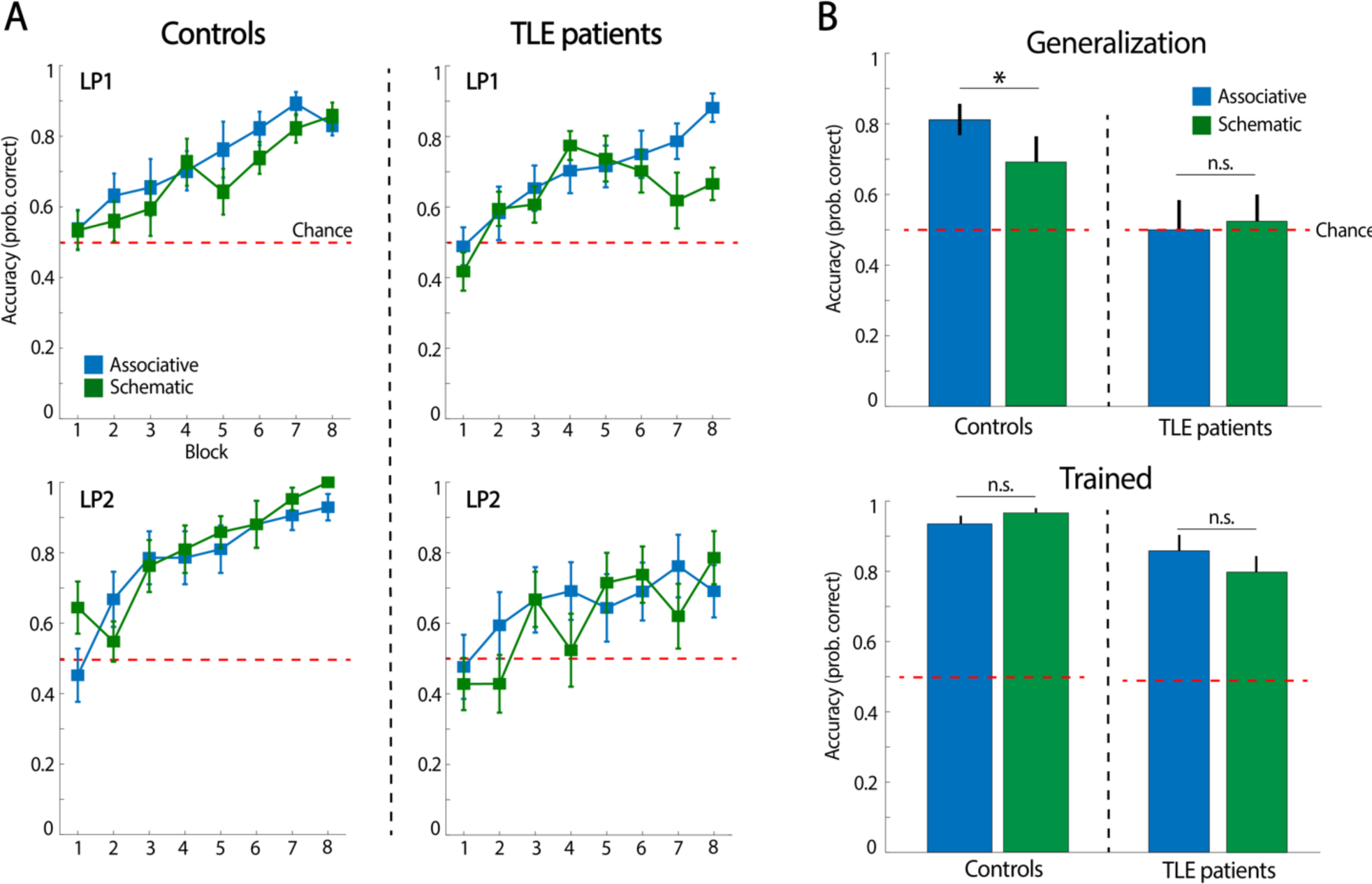
Behavioural data in TLE and healthy control participants (Experiment 2). (**A**) Averaged participants’ accuracy in selecting the correct scene association with a given face throughout LP1 and LP2 for each experimental condition. (**B**) Averaged participants’ accuracy in the generalization and trained memory tests. *p < 0.05 and n.s., p > 0.05. Error bars indicate standard error of the mean.

We next searched for group differences in the generalization test through an ANOVA, including condition and group as a within- and between-subject factors, respectively. This analysis revealed a significant main effect of group (F (1, 26) = 9.22, p = 0.005) but not of condition (F (1, 26) = 0.59, p = 0.45), nor a condition × group interaction (F (1, 26) = 1.30, p = 0.26) (Figure 5B), thereby indicating that TLE patients showed poorer ability to generalize in the associative and schematic condition. In fact, while behavioural accuracy in each of the conditions was high and above chance in healthy controls (Associative: t(13) = 6.87; p < 0.01; Schematic: t(13) = 2.57; p = 0.02), TLE patients performed at chance in the tests (Associative: t(13) = 0.31; p = 0.76; Schematic: t(13) = 0.30; p = 0.76). A paired-sample *t*-test analysis showed that accuracy did not differ statistically between conditions at within group level (controls: t (13) = 1.82, p = 0.09; TLE patients: t (13) < 0.5).

Finally, regarding memory accuracy for trained events, an ANOVA including condition and group as a within- and between-subject factors, respectively, showed a significant group (F (1, 26) = 13.61, p = 0.01) but not a group × condition interaction effect (F (1, 26) = 2.13, p = 0.16), indicating that TLE patients were less accurate in recognising the individual face-scene associations acquired during LP2 when compared to controls (Figure 5B). Importantly, both controls and TLE patients showed consistent above-chance performance in each of the test measures (controls – associative condition: t(13) = 14.14; p < 0.01; controls – schematic: t(13) = 20.21; p < 0.01; TLE patients – associative: t(13) = 1.94; p = 0.07; TLE patients – schematic: t(13) = 5.1; p < 0.01). A paired-sample *t*-test analysis showed that accuracy did not differ statistically between conditions at within-group level (controls: t (13) = −0.56, p = 0.58; TLE patients: t (13) = −1.33, p = 0.21).

## Discussion

A challenge in memory research has been to understand how structures of knowledge can aid integrative encoding of new information. Here, we showed that process is hippocampus-dependent, as TLE patients were impaired in inferring novel relations between elements within mnemonic networks but not in retrieving individual pictures. In addition, we observed more persistent and widespread theta activity (3-6 Hz) in the scalp during the encoding of novel pictures related to schematic memory networks, suggesting that theta may reflect distances between elements within an arbitrary representational network space. Finally, we found high similarity values for neural activity patterns elicited by novel and related events only within associative networks, thereby suggesting that neural reactivation may be important in integrative encoding only when novel information relates to mnemonic network of elements linked by direct associations.

Our findings provide evidence that MTL structures, including the hippocampus, are essential to enabling the rapid integration of new information within stored knowledge in a schematic structure. These results align well with previous studies that revealed the critical role of the hippocampus in enabling inferential learning between different episodic events that overlap in the perceptual content in healthy individuals (Schlichting and Preston 2015; Zeithamova et al., 2012); and in patients with lesions in the MTL (Pajkert et al. 2017). However, the current findings extend these findings by showing that the hippocampus-dependent nature of this process also affects inferential learning that relies on relational links between distinct episodes whose overlapping content is at the conceptual level (i.e., stimulus category). The degree to which hippocampal integration mechanisms identified in episodic inference contribute to other forms of generalization, such as concept learning, has often been neglected in the literature. However, recent fMRI findings in humans showed that the anterior hippocampus, in concert with the PFC, generated and tracked the prototype representation of multiple items that overlapped in their content, and the degree to which participants relied on such conceptual representation abstracted across the training set predicted their ability to generalize in a later test (Bowman and Zeithamova 2018). In addition, the notion that the hippocampus is critical in enabling the integration of new items within a schematic network has strong support from animal and human studies showing that the existence of prior knowledge promoted the rapid assimilation of new but related information (van Kesteren et al. 2010; Packard et al. 2017; Tse et al. 2007, 2011) into hierarchically organized memory networks in the hippocampus (McKenzie et al. 2014).

We observed that the learning-eliciting theta oscillations during the integration of novel information into an existing memory network reflected its underlying organizational properties. Specifically, we found that theta activity persisted longer when new information was linked to schematic memory networks than when novel items had to be integrated into memory networks with an associative structure. These findings suggest that theta oscillations is a putative neural mechanism by which our brain searches for memories throughout the representational space. Indeed, a plethora of animal (Buzsáki 2002; Buzsáki, Lai-Wo S., and Vanderwolf 1983) and human studies have revealed the relevant role of theta oscillations in supporting spatial memory and navigation (Ekstrom et al. 2005; Jacobs et al. 2013).Several recent studies have suggested that theta power may itself correlate with spatial distances (Bush et al. 2017; Vass et al. 2016). Intriguingly, a recent study using deep electrodes in the hippocampus of human epileptic patients showed that theta activity coded semantic distances between words from a list (Solomon et al. 2019), thereby lending support to the notion that theta oscillations reflect relations between nonspatial items in memory. Our findings that theta elicited greater response from events related to schematic memory networks than those related to associative networks contribute to the idea that theta oscillations may signal memory trajectories in the representational space. These findings contribute to the idea that theta oscillations may signal memory trajectories in a general-domain representational space.

Prior literature has emphasized how inferential learning relies on mechanisms of memory reactivation (e.g., (Zeithamova et al., 2012)). These studies revealed that prior event details are reinstated at the encoding of related experiences and that this supports participants’ ability to infer relationships between distinct events that share content. The current study builds upon and significantly extends prior studies by showing that the representational nature and temporal dynamics of such reactivation during inferential encoding depend on the organizational structure of the related memory network. Indeed, our representational similarity analysis revealed high correlation between EEG patterns elicited by events encoded in LP2 and EEG patterns triggered by related events from the previous LP1 phase. However, while we found a similarity increase in a large cluster of time points, from ~500 to 2500 ms stimulus onset, between EEG-elicited patterns with new and related events within the associative network, neural similarity between new events and events within the related schematic network was only higher for a brief window of time. This differential pattern of similarity results suggests that the reactivation nature of prior memory events during integrative encoding may adapt as a function of the structural properties of the related memory network. Accordingly, only novel events that largely overlap in content with episodic content from an associative memory network would entail a detailed reactivation of the related event. We reasoned that such an adaptive property of the memory systems would be optimal in our daily life activity, as most of our experienced events ultimately share a relationship with stored memories. We conjecture that an efficient regulatory mechanism may exist to avoid the costs of inducing memory reactivation of specific memories during the ongoing encoding, while maximizing the benefits of maintaining a linked memory representation of an encoded event with previously stored memory representations. We speculate that this regulatory mechanism may be guided by the ability of a reminder/cue to navigate throughout the representational memory space and rapidly find associated events. If they are found, then these memory representations are reactivated and integrated in a common representational space (i.e., associative network in the current experiment). However, when a given cue does not match a specific representation for an event, then it moves to higher levels of representation, such as the conceptual level, thereby maximizing the ability to keep the current encoded event related to other memories that overlap at this representational level, as in the schematic network condition in the present experiment. Intriguingly, the pattern of similarity results that we obtained in the schematic condition fit recent EEG findings in humans revealing that information flow during encoding and retrieval may be reversed in order (Linde-Domingo et al. 2019). Thus, while visual encoding starts with low-level perceptual followed by high-level abstract processing, the mnemonic stream can prioritize the access to conceptual information. Our finding that the EEG patterns from early temporal windows elicited by novel events were similar to EEG patterns elicited at a late temporal window by the related event within a schematic memory network suggests that a similar retrieval-oriented prioritization to access conceptual information may be engaged in the schematic condition in our study.

Taken together, our findings shed light on the neural mechanisms that support the integration of novel information into existing memory networks. Our central finding is that both associative and schematic structures of memory networks aid integrative encoding of new information via the hippocampus, but that they engage different theta and neural reactivation patterns. Theta oscillatory activity was more persistent and widespread when the encoded event was to be integrated into a schematic memory network, lending support to the notion that theta oscillations may reflect distances between elements within representational network space. On the other hand, a stronger and temporally extended reactivation of prior event memories was found only during the encoding of events that were integrated into associative memory networks, thereby suggesting the existence of regulatory mechanisms that promote the reactivation of related memory events when they belong to an associative structure in which multiple events are linked by direct association within a network. More broadly, the results emphasize the flexible nature of memory, whereby novel experiences and organizational properties of stored knowledge interact to enable structured representation of information in an ever-changing environment.

## REFERENCES

Backus, Alexander R. et al. 2016. “Hippocampal-Prefrontal Theta Oscillations Support Memory Integration.” Current Biology 26(4): 450–57.

Blair, R.Clifford, and Walt Karniski. 1993. “An Alternative Method for Significance Testing of Waveform Difference Potentials.” Psychophysiology 30(5): 518–24. http://doi.wiley.com/10.1111/j.1469-8986.1993.tb02075.x (December 8, 2019).

Bowman, Caitlin R., and Dagmar Zeithamova. 2018. “Abstract Memory Representations in the Ventromedial Prefrontal Cortex and Hippocampus Support Concept Generalization.” Journal of Neuroscience 38(10): 2605–14.

Bush, Daniel et al. 2017. “Human Hippocampal Theta Power Indicates Movement Onset and Distance Travelled.” Proceedings of the National Academy of Sciences of the United States of America 114(46): 12297–302.

Buzsáki, György. 2002. “Theta Oscillations in the Hippocampus.” Neuron 33(3): 325–40.

Buzsáki, György, Leung Lai-Wo S., and Cornelius H. Vanderwolf. 1983. “Cellular Bases of Hippocampal EEG in the Behaving Rat.” Brain Research Reviews 6(2): 139–71.

Delorme, Arnaud, and Scott Makeig. 2004. “EEGLAB: An Open Source Toolbox for Analysis of Single-Trial EEG Dynamics Including Independent Component Analysis.” Journal of Neuroscience Methods 134(1): 9–21.

Dusek, Jeffery A., and Howard Eichenbaum. 1997. “The Hippocampus and Memory for Orderly Stimulus Relations.” Proceedings of the National Academy of Sciences of the United States of America 94(13): 7109–14.

Eichenbaum, Howard. 2017. “Memory: Organization and Control.” Annual Review of Psychology 68(1): 19–45.

Ekstrom, Arne D. et al. 2005. “Human Hippocampal Theta Activity during Virtual Navigation.” Hippocampus 15(7): 881–89.

Greene, Anthony J., William L. Gross, Catherine L. Elsinger, and Stephen M. Rao. 2006. “An FMRI Analysis of the Human Hippocampus: Inference, Context, and Task Awareness.” Journal of Cognitive Neuroscience 18(7): 1156–73.

Groppe, David M., Thomas P. Urbach, and Marta Kutas. 2011. “Mass Univariate Analysis of Event-Related Brain Potentials/Fields I: A Critical Tutorial Review.” Psychophysiology 48(12): 1711–25.

Heckers, Stephan et al. 2004. “Hippocampal Activation during Transitive Inference in Humans.” Hippocampus 14(2): 153–62.

Jacobs, Joshua et al. 2013. “Direct Recordings of Grid-like Neuronal Activity in Human Spatial Navigation.” Nature Neuroscience 16(9): 1188–90.

Van Kesteren, Marlieke T.R., Guillén Fernández, David G. Norris, and Erno J. Hermans. 2010. “Persistent Schema-Dependent Hippocampal-Neocortical Connectivity during Memory Encoding and Postencoding Rest in Humans.” Proceedings of the National Academy of Sciences of the United States of America 107(16): 7550–55.

Van Kesteren, Marlieke T.R., Dirk J. Ruiter, Guillén Fernández, and Richard N. Henson. 2012. “How Schema and Novelty Augment Memory Formation.” Trends in Neurosciences 35(4): 211–19.

van Kesteren, Marlieke T R, Guillén Fernández, David G Norris, and Erno J Hermans. 2010. “Persistent Schema-Dependent Hippocampal-Neocortical Connectivity during Memory Encoding and Postencoding Rest in Humans.” Proceedings of the National Academy of Sciences of the United States of America 107(16): 7550–55. http://www.ncbi.nlm.nih.gov/pubmed/20363957 (December 8, 2019).

Langner, Oliver et al. 2010. “Presentation and Validation of the Radboud Faces Database.” Cognition and Emotion 24(8): 1377–88.

Linde-Domingo, Juan, Matthias S. Treder, Casper Kerrén, and Maria Wimber. 2019. “Evidence That Neural Information Flow Is Reversed between Object Perception and Object Reconstruction from Memory.” Nature Communications 10(1): 179. http://www.nature.com/articles/s41467-018-08080-2 (December 8, 2019).

Maris, Eric, and Robert Oostenveld. 2007. “Nonparametric Statistical Testing of EEG- and MEG-Data.” Journal of Neuroscience Methods 164(1): 177–90.

McKenzie, Sam et al. 2014. “Hippocampal Representation of Related and Opposing Memories Develop within Distinct, Hierarchically Organized Neural Schemas.” Neuron 83(1): 202–15.

Minear, Meredith, and Denise C. Park. 2004. “A Lifespan Database of Adult Facial Stimuli.” Behavior Research Methods, Instruments, and Computers 36(4): 630–33.

Packard, Pau A. et al. 2017. “Semantic Congruence Accelerates the Onset of the Neural Signals of Successful Memory Encoding.” Journal of Neuroscience 37(2): 291–301.

Pajkert, Anna et al. 2017. “Memory Integration in Humans with Hippocampal Lesions.” Hippocampus 27(12): 1230–38.

Preston, Alison R., Yael Shrager, Nicole M. Dudukovic, and John D.E. Gabrieli. 2004. “Hippocampal Contribution to the Novel Use of Relational Information in Declarative Memory.” Hippocampus 14(2): 148–52.

Sans-Dublanc, A., E. Mas-Herrero, J. Marco-Pallarés, and L. Fuentemilla. 2017. “Distinct Neurophysiological Mechanisms Support the Online Formation of Individual and Across-Episode Memory Representations.” Cerebral Cortex 27(9): 4314–25.

Schlichting, Margaret L., and Alison R. Preston. 2015. “Memory Integration: Neural Mechanisms and Implications for Behavior.” Current Opinion in Behavioral Sciences 1: 1–8.

Schlichting, Margaret L., and Alison R. Preston. 2016. “Hippocampal–Medial Prefrontal Circuit Supports Memory Updating during Learning and Post-Encoding Rest.” Neurobiology of Learning and Memory 134(Part A): 91–106. http://dx.doi.org/10.1016/j.nlm.2015.11.005.

Silva, Marta, Christopher Baldassano, and Lluís Fuentemilla. 2019. “Rapid Memory Reactivation at Movie Event Boundaries Promotes Episodic Encoding.” The Journal of neuroscience: the official journal of the Society for Neuroscience 39(43): 8538–48.

Solomon, Ethan A., Bradley C. Lega, Michael R. Sperling, and Michael J. Kahana. 2019. “Hippocampal Theta Codes for Distances in Semantic and Temporal Spaces.” Proceedings of the National Academy of Sciences 116(48): 24343–52. http://www.pnas.org/lookup/doi/10.1073/pnas.1906729116 (December 8, 2019).

Sols, Ignasi, Sarah Dubrow, Lila Davachi, and Lluís Fuentemilla Correspondence. 2017. “Event Boundaries Trigger Rapid Memory Reinstatement of the Prior Events to Promote Their Representation in Long-Term Memory.” Current Biology 27: 3499–3504. https://doi.org/10.1016/j.cub.2017.09.057 (December 8, 2019).

Tse, Dorothy et al. 2007. “Schemas and Memory Consolidation.” Science 316(5821): 76–82.

Tse, Dorothy et al. 2011. “Schema-Dependent Gene Activation.” Science 333(August): 891–95.

Vass, Lindsay K. et al. 2016. “Oscillations Go the Distance: Low-Frequency Human Hippocampal Oscillations Code Spatial Distance in the Absence of Sensory Cues during Teleportation.” Neuron 89(6): 1180–86.

Watrous, Andrew J. et al. 2013. “Frequency-Specific Network Connectivity Increases Underlie Accurate Spatiotemporal Memory Retrieval.” Nature Neuroscience 16(3): 349–56.

Xiao, Jianxiong et al. 2010. “SUN Database: Large-Scale Scene Recognition from Abbey to Zoo.” In Proceedings of the IEEE Computer Society Conference on Computer Vision and Pattern Recognition,, 3485–92.

Zeithamova, Dagmar, April L. Dominick, and Alison R. Preston. 2012. “Hippocampal and Ventral Medial Prefrontal Activation during Retrieval-Mediated Learning Supports Novel Inference.” Neuron 75(1): 168–79.

Zeithamova, Dagmar, and Alison R. Preston. 2010. “Flexible Memories: Differential Roles for Medial Temporal Lobe and Prefrontal Cortex in Cross-Episode Binding.” Journal of Neuroscience 30(44): 14676–84.

Zeithamova, Dagmar, Margaret L. Schlichting, and Alison R. Preston. 2012. “The Hippocampus and Inferential Reasoning: Building Memories to Navigate Future Decisions.” Frontiers in Human Neuroscience 6(March 2012): 1–14.

